# Salt-induced reduction of hyperswarming motility in *Bacillus cereus* MHS is associated with reduced flagellation, reduced nanotube formation and reduction in expression of quorum sensing regulator

**DOI:** 10.1101/2024.12.17.628887

**Authors:** Nirbhay Kumar Bhadani, Hemanta Sarmah, Kritika Prasad, Nisha Gupta, Tapas K Sengupta

## Abstract

Bacteria have been known to thrive in challenging environmental niches through diverse phenomena. Swarming is one such favourable adaptation that could help bacteria survive extreme conditions. Therefore, targeting swarming is crucial for improving our understanding of bacterial motility and preventing related infections. *Bacillus cereus,* which causes food poisoning, has been shown to perform swarming, and salts like NaCl can act as a food preservative to control bacterial growth. In order to explore the possible alterations in the swarming of *Bacillus cereus* in the presence of salt, the present study encompasses the effect of NaCl on the swarming characteristics of a natural bacterial isolate, *Bacillus cereus* MHS, with a hyperswarming phenotype. Here we report that the presence of an increased amount of NaCl in growth media could induce a reduction in swarming motility and swarming pattern of MHS on Luria agar plates and the observed reduction in swarming was associated with the reduced flagellation and reduction in bacterial nanotube formation. Gene expression studies supported the phenotypic and ultrastructure observations as the expressions of *bfla* and *ymdb* genes, involved in formations of flagella and nanotubes respectively, were found to be reduced in the swarming MHS cells in the presence of increased NaCl in swarm media. It was also observed that the salt-induced reduction in swarming of MHS is associated with the reduced expression of the quorum sensing regulator gene *plcR.* This study first time reports the bacterial nanotubes in a *Bacillus cereus* strain and indicates a possible link between the bacterial nanotube formation and hyperswarming phenotype in *Bacillus cereus* MHS.

## 1. Introduction

Swarming refers to the movement of individuals in a group and has been commonly observed in nature such as in schools of fish, flocks of birds, and also in several bacterial species. Swarming in bacteria is defined as a collective and coordinated movement of flagellated bacterial cells over surfaces under certain specific conditions (Kearns, 2010). Group translocation as seen in swarming, provides a competitive advantage over a single individual and helps to move faster for exploring food resources and can also act as a defense mechanism to evade predators. Thus, a social behaviour like swarming is considered advantageous for bacterial growth, overcoming stresses, and adapting to adverse environmental cues. In laboratory conditions, the swarming behaviour depends on various factors like water content, agar concentration, nutrient availability or composition, incubation temperature, etc. (Rashid & Kornberg, 2000; Tremblay & Déziel, 2008; Tremblay et al., 2007).

*Bacillus cereus* is a gram-positive, rod-shaped, motile, spore-forming bacteria. *B. cereus* causes emetic syndrome through cereulide toxins encoded by plasmid-borne cereulide synthetase genes. In contrast, diarrhoea is caused by the ingestion of vegetative cells or spores, producing enterotoxins like hemolysin B, nonhemolytic enterotoxin, and cytotoxin K in the small intestine (Dietrich et al., 2021). Besides food poisoning, *B. cereus* can cause other infections like bacteremia, endophthalmitis, and gas gangrene-like cutaneous infections in immunocompromised and immunocompetent individuals, intravenous drug abusers, neonates, and individuals with catheters (Bottone 2010).

*Bacillus cereus* has often been considered a menace to the food industry for its role in food spoilage, and decreased food quality and safety (Salvetti et al. 2011). Moreover, *Bacillus cereus* has been reported to swarm over surfaces mainly depending on nutrient availability (Liu et al., 2020) which helps it to colonize surfaces.

Sodium chloride is used as a food preservative and acts by depriving the bacteria of using available water as a nutrient. The growth of bacteria is avoided or delayed in the presence of salts (Elias et al., 2020). Since different foods get generally contaminated on the surface (Stecchini et al., 2000), the effect of salt (NaCl) on the swarming of *Bacillus cereus* is an important aspect to study.

In the present study, we report a salt tolerant *Bacillus cereus* MHS strain that was isolated from a pond and found to be a natural hyperswarmer. In our experiments, we regulated the hyperswarming ability of this strain by increasing NaCl concentrations in the growth media to study the changes in swarming, morphology, and level of flagellation of *Bacillus cereus* MHS. Swarming requires communication among bacteria which is achieved by quorum sensing mechanisms *via* signaling molecules like autoinducers and signaling peptides. Bacteria also communicate via direct contact, including genetic material exchange through conjugation and pilus formation (Lederberg & Tatum, 1946). Additionally, structures like nanowires in *Shewanella oneidensis* MR-1 facilitate electron transport (Pirbadian et al., 2014), while nanopods in *Delftia sp. Cs1-4* acts as a conduit from the cells composed of outer membrane vesicles with a surface layered protein npdA (Shetty et al., 2011). Moreover, Dubey and Yehuda (2011), demonstrated the presence of another type of bacterial communication mediated by nanotubes in *Bacillus subtilis* that serve as a pathway for the molecular exchange of hereditary as well as non-hereditary cargos, delivery of toxic substances to competitor neighbouring cells as a strategy to fight against it or take away nutrients from neighbouring bacteria which were found to be abolished when nanotubes were perturbed. The presence of nanotubes and the role of genes involved in their regulation have not been deciphered yet in *Bacillus cereus*. Therefore, an attempt was made to report the presence of bacterial nanotubes for the first time in a naturally occurring hyperswaming *Bacillus cereus* MHS strain and its alterations on increasing NaCl concentrations. Moreover, we also studied the expression of genes like *bfla* for motility, *plcr* for quorum sensing during swarming under increasing NaCl concentrations. Since *ymdb* which encodes calcineurin like phosphodiesterase has been reported to be involved in the formation of nanotubes in *Bacillus subtilis* (Dubey & Ben-Yehuda, 2011), we studied the expression of *ymdb* for nanotube formation in our strain and speculate a role of *ymdb* in hyperswarming of *Bacillus cereus* MHS.

## 2. Materials and Methods

### 2.1. Bacterial strains and phylogenetic analysis

*Bacillus cereus* strain MHS used in the present study was isolated from a water sample collected from a pond (22°57’15.9’’N 88°31’20.9” E) within the IISER Kolkata campus and no permission was necessary for sample collection. The strain MHS was grown and routinely maintained in Luria-Bertini broth (LB) (10g L^-1^ NaCl, 10g L^-1^ tryptone, 5g L^-1^ yeast extract) with shaking at 150 rpm at 30°C and in semi-solid Luria agar (10g L^-1^ NaCl, 10g L^-1^ tryptone, 5g L^-1^ yeast extract, 0.7% agar). The bacterial strain was stored in glycerol stocks (15% glycerol) for future use.

For *Bacillus cereus* MHS, sequencing of the *16S rDNA* gene was done to identify the strain lineage from the known sequences in NCBI (Altschul et al. 1990). A pure colony of MHS was grown in LB medium and incubated at 30°C under shaking conditions (150rpm). 1.5 mL of grown bacterial culture was centrifuged at 10000rpm for 3 minutes.

The genomic DNA of bacteria was isolated using HiPurA^TM^ Bacterial Genomic DNA purification kit (HiMedia). The isolated genomic DNA was used as a template to amplify *16s rDNA* using 27F and 1492R universal primers (Table 1) (Dasgupta et al., 2013). The amplified gene was analyzed by agarose gel electrophoresis. DNA purification from agarose gel was performed using PureLink^TM^ Quick Gel Extraction Kit (Invitrogen) as per the manufacturer’s manual. All the required buffers were provided by the kit. The amplified gene (bands) sequences were determined from the purified PCR products using genetic analyzer 3510 (Applied Biosystems) with Big Dye^TM^ Terminator Cycle Sequencing kit v3.1.

**Table 1:**
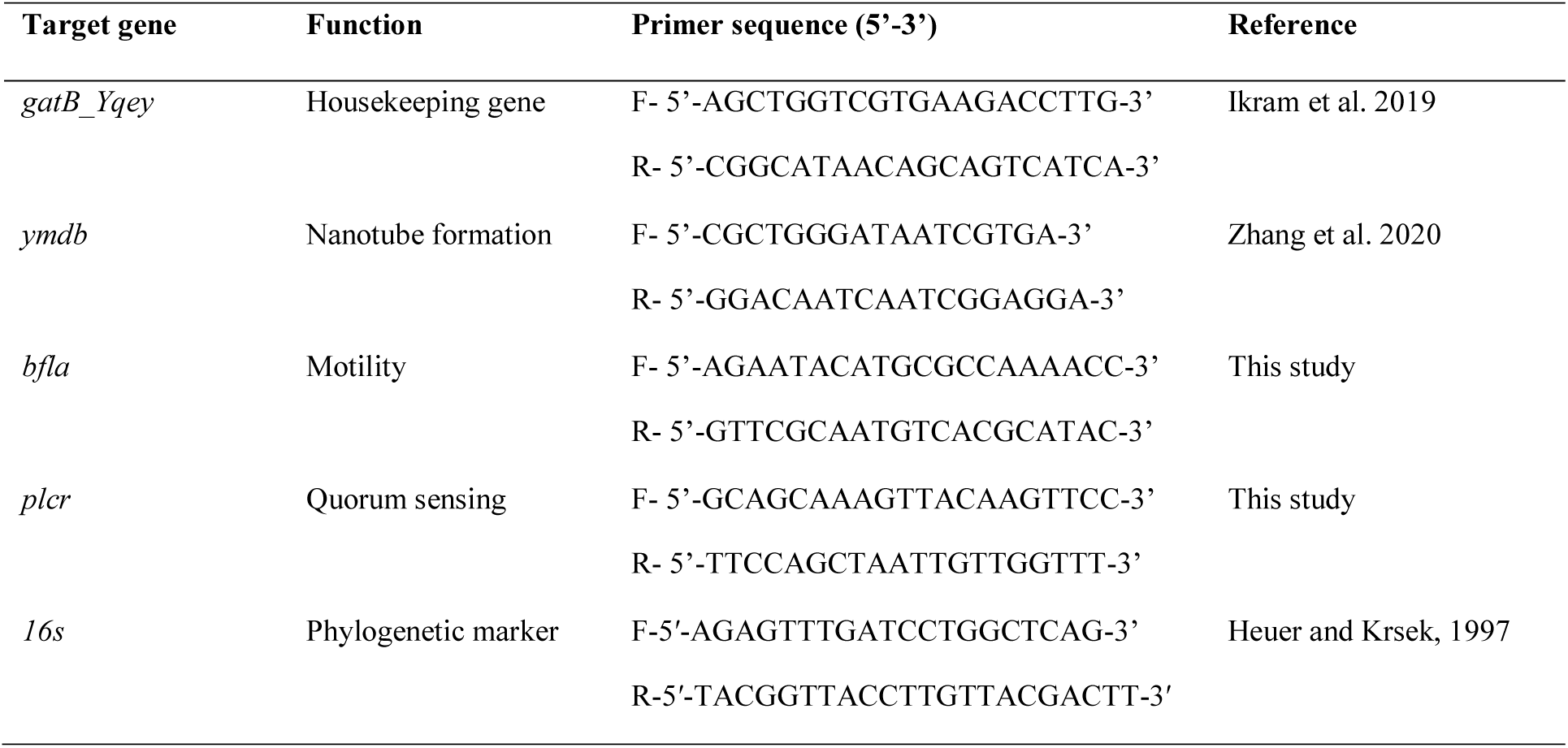
List of primers used for semi-quantitative PCR.

The *16S rDNA* gene sequence of MHS was deposited in the National Center for Biotechnology Information (NCBI, accession number: KR132556). The sequence of *16S rDNA* of MHS was compared with the known sequences using the Basic Local Alignment Sequencing Tool (BLAST) of the National Institute of Health (NIH, Bethesda, Maryland, USA). Twenty-eight sequences of the cultivable bacteria were screened from BLAST search, and a multiple sequence alignment was done using CLUSTALX2. The alignment was scrutinized through SeaView for errors and was further edited accordingly. Using MEGA v4.0, a phylogenetic tree was constructed by the neighbour-joining method, Kimura-2-parameter. The tree was consequently bootstrapped at 1000 replicates (Dasgupta et al., 2015).

In the present study, the swarming ability of the *Bacillus cereus* MHS strain was compared to the relative swarming of a normal swarming *Bacillus cereus* strain MSM-S1 (Chakraborty et al., 2016).

### 2.2. Bacterial growth and swarming motility assay

To monitor the growth of *B. cereus* MHS in LB media in the presence of different NaCl concentrations (1.0, 2.0, and 3.0%), 100µL of overnight grown bacterial culture (10^7^ cfu/mL) was inoculated in LB in the absence or presence of NaCl and incubated under shaking conditions (150 rpm) at 30°C. Then the optical density (O.D.) of the culture was measured at 600nm at an interval of 1 hour using a UV-Vis spectrophotometer (Shimadzu UV-1800). The O.D. values were plotted against the incubation time to obtain the bacterial growth curve (Chakraborty et al., 2016).

Experiments to study the swarming ability of *B. cereus* MHS under different NaCl concentrations (1.0, 2.0, and 3.0%) were done on 0.7% LA plates. For that, *B. cereus* MHS from a fresh LA plate was inoculated in 5 mL of LB as a seed culture and grown overnight at 30°C under shaking conditions (150 rpm). 2.5µL of overnight grown cell suspension (adjusted to have 10^7^ cfu/mL based on O.D. at 600 nm) was seeded by spot inoculation onto the center of LA plates and incubated at 30^°^C for 12 h. The images of growing colonies were taken by using a gel documentation system (Syngene G: Box). The area covered by the colonies after 12 hours was calculated by the ImageJ software 1.54g.

### 2.3. Confocal laser scanning microscopy

For observing the morphology, orientation, and organization of cells in colonies of *B. cereus* MHS during swarming under different NaCl (1.0, 2.0, and 3.0%) concentrations, confocal laser scanning microscopy (CLSM) was performed. The peripheral regions of growing bacterial colonies (along with the agar base) were cut into cubes and placed onto glass-bottom dishes (30mm, SPL Life Sciences). They were then stained with 0.001% acridine orange (w/v) and incubated for 20 minutes in the dark, followed by gentle washing twice with 100µL of phosphate-buffered saline (PBS), pH 7. The cells in the intact colony edge were visualized under CLSM-710 (Axio Observer Microscope, Version Z.1; Carl Zeiss, Germany) with argon 488nm laser line, and corresponding emission of 493 to 576 nm was recorded. Throughout the experiment, images of 8-bit depth having a frame size of 512X512 were maintained. The length, width, and surface area to volume ratio of the bacterial cells were calculated manually using ImageJ (Chakraborty et al., 2016).

### 2.4. Transmission electron microscopy

Transmission electron microscopy (TEM) was performed to observe the presence and abundance of flagella in the *B. cereus* MHS cells during swarming under different NaCl concentrations (1.0, 2.0, and 3.0%). For that, cells were taken from the peripheries of growing colonies, resuspended in PBS and 5 µL of this bacterial suspension was placed on the formvar-coated copper grid (300 mesh size, ted Pella, INC). Cells were fixed in 2.0% paraformaldehyde for 1 minute and excess liquid was wicked off without completely drying the grid to prevent flagellar shearing. The grids were then negatively stained with 2.0% (w/v) phosphotungstic acid for 3 minutes followed by air-drying. Fixed and stained cells were viewed under a transmission electron microscope at 120 KV (JEOL, JEM-2100 plus electron microscope) (Roy et al., 2010).

### 2.5. Field emission scanning electron microscopy

To obtain a better resolution of the surface topography of *B. cereus* MHS during swarming on 0.7% LA under different NaCl concentrations (1.0, 2.0, and 3.0 %), Field Emission Scanning Electron Microscopy (FESEM) was performed. Cells from the periphery of growing colonies were collected, smeared onto clean coverslips, and washed twice with PBS. Next, the coverslips were flooded with 2.5% glutaraldehyde (made in PBS) and incubated for 2 hours at room temperature. After incubation, the samples were washed thrice for 5 minutes each time with PBS and 0.1% osmium tetroxide (made in PBS) was added on top of the samples followed by incubating in the dark for 40 minutes. After incubation, samples were washed thrice with PBS. Cells were gradually dehydrated using 30%, 40%, 50%, 60%, 70%, 80%, and 90% ethanol for 10 minutes and finally with 100% ethanol for 15 minutes. Next, the coverslips were air-dried overnight and further allowed to dry in a desiccator and taken for sampling the following day. Before imaging, the coverslips with the samples on them were coated with a thin layer of conducting metal, specifically by gold-palladium in a sputter coater. Finally, cells were observed under Supra 55 VP, field emission scanning electron microscope (Carl Zeiss) using SmartSEM software and images were taken (Chakraborty et al., 2016).

### 2.6. Semi-quantitative reverse transcription-polymerase chain reaction

To check the expression of genes related to flagella formation (*bfla*), nanotube formation (*ymdb*), and quorum sensing (*plcr*) in *Bacillus cereus* MHS during swarming on LA plate under different NaCl concentrations (1.0, 2.0, and 3.0%), semi-quantitative reverse transcription-polymerase chain reaction (RT-PCR) was done. For that, peripheral cells were collected from the growing colonies and the total RNA was isolated by the protocol described by (Oh & So, 2003) with some modifications. Cells were ruptured using glass beads (425-600 µm in diameter, Sigma) and RNAiso Plus (Takara) was added to the ruptured cells before the isolation of RNA. To remove DNA contamination, the isolated RNA was further treated with DNase I (Thermo Fisher Scientific). Purified RNA was quantified using a Nanodrop spectrophotometer (BioTek Epoch Microplate Spectrophotometer, Agilent Technologies Inc). The RNA (500 ng) was then reverse-transcribed using the PrimeScript 1st strand cDNA synthesis kit (Takara) following the instructions provided by the manufacturer. The prepared cDNA was used as a template for PCR by the primer sets of genes mentioned in Table 1. Semi-quantitative reverse transcription PCR was performed for *gatB_Yqey and ymdb* for 25 and 32 cycles, respectively, while for *bfla,* and *plcr* genes, PCR was done for 35 cycles. The amplified PCR products were analyzed by agarose gel electrophoresis. The intensity of the amplified PCR product bands was measured using the GeneTools software. Differential expression of genes was measured by normalizing the band intensity of a gene with the band intensity of the housekeeping gene (*gatB_Yqey*) and measuring the fold change.

### 2.7. Statistical analysis

All the above experiments were performed independently at least three times and the data was expressed as mean ± SE. Microsoft Excel and GraphPad Prism 6 were used to analyze the data. Student’s *t*-test (2-tailed, unpaired) was used to analyze the statistical significance. *P-value* lesser than 0.05 was considered to be statistically significant (\**P-value* ≤ 0.05; \*\**P-value* ≤ 0.01; \*\*\**P* value ≤ 0.001; ****, *P-value* ≤ 0.0001).

## 3. Results

### 3.1. Bacterial characterization and phylogenetic analysis

The bacterial isolate’s 16S rDNA sequence was compared with the available sequences in GenBank with the BLAST tool. The phylogenetic analysis revealed that the isolated bacterial strain was closely related to *Bacillus cereus* (**Figure S1**) and thus named *Bacillus cereus* MHS. The strain was found to be highly motile because when spotted on a 0.7% LA plate, the colony of MHS could cover more than half of the 90 mm petri dish within 12h and hence this strain was termed hyperswarmer bacteria. The other strain, *Bacillus cereus* MSM-S1, could not cover the plate as fast as *Bacillus cereus* MHS (**Figure 1**), hence it was termed normal swarming bacteria.

**Fig. 1:**
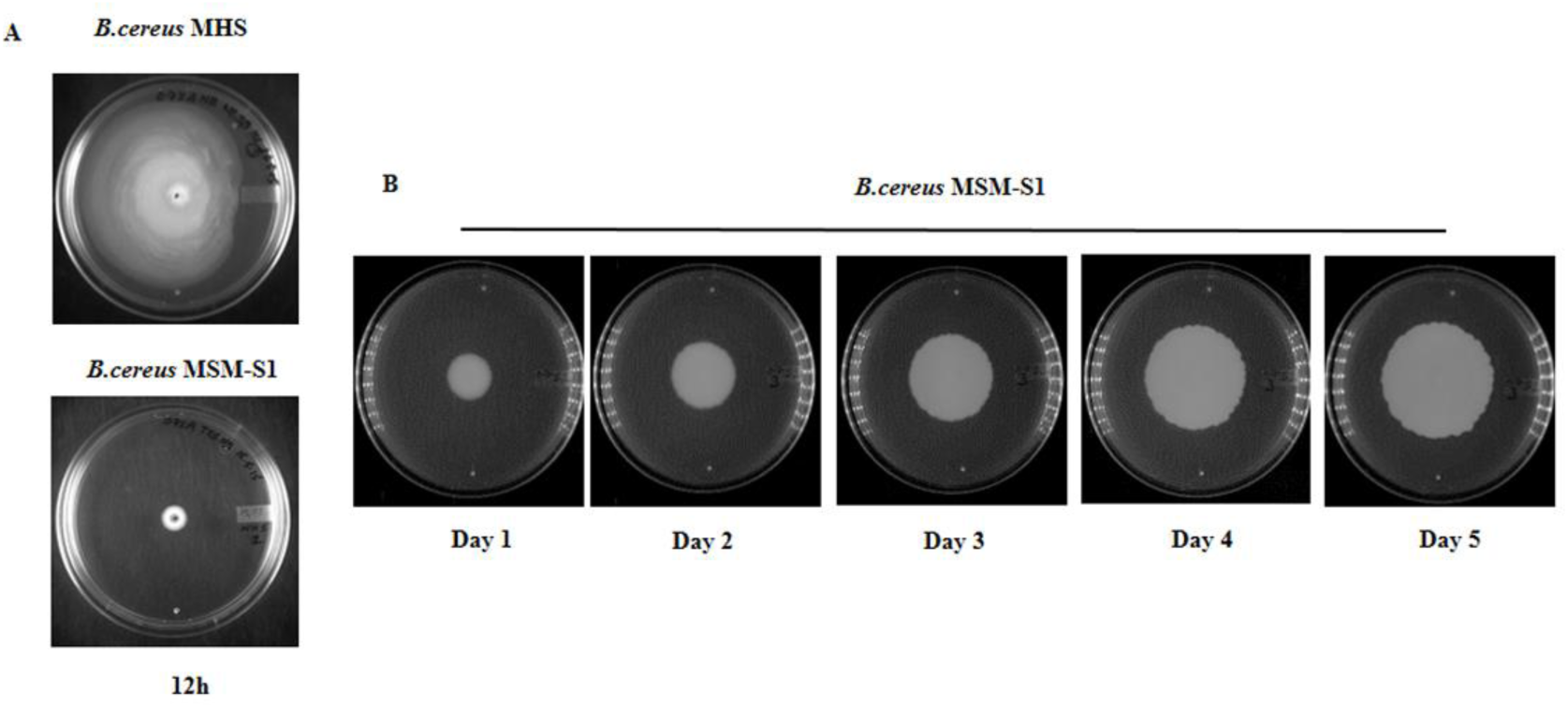
Representative images of (A) Swarming motility of *Bacillus cereus* MHS and MSM-S1 after 12h of incubation in the 0.7% LA plate; (B) Swarming motility of *Bacillus cereus* MSM-S1 in 0.7% LA plate for 5 days

### 3.2. Effect of differential salt concentrations on growth and swarming motility of Bacillus cereus MHS

The growth and swarming motility of *Bacillus cereus* MHS was observed to be significantly reduced with increasing salt concentrations in a dose-dependent manner. An increase in salt concentrations from 1.0% to 3.0% affected the growth of *Bacillus cereus* MHS cells and resulted in an increase in lag periods and generation times of MHS when grown in liquid media **(Figure 2 A)**. A similar effect of salt on the swarming motility of the *Bacillus cereus* MHS colony was observed when grown on 0.7% LA plates **(Figure 2 B)**. In the presence of 1.0% NaCl, the area covered by the *Bacillus cereus* MHS colony was the maximum (4014 ± 458.7 mm^2^) and it was also almost more than half the available plate areas after 12h. This is typically a hyperswarming phenotype. In the presence of 2.0% NaCl, the area covered by the swarming colonies was intermediate (648.1 ± 84.7 mm^2^) while it was minimum in the presence of 3.0% NaCl (52.5 ± 5.2 mm^2^) **(Figure 2 C).**

**Fig. 2.**
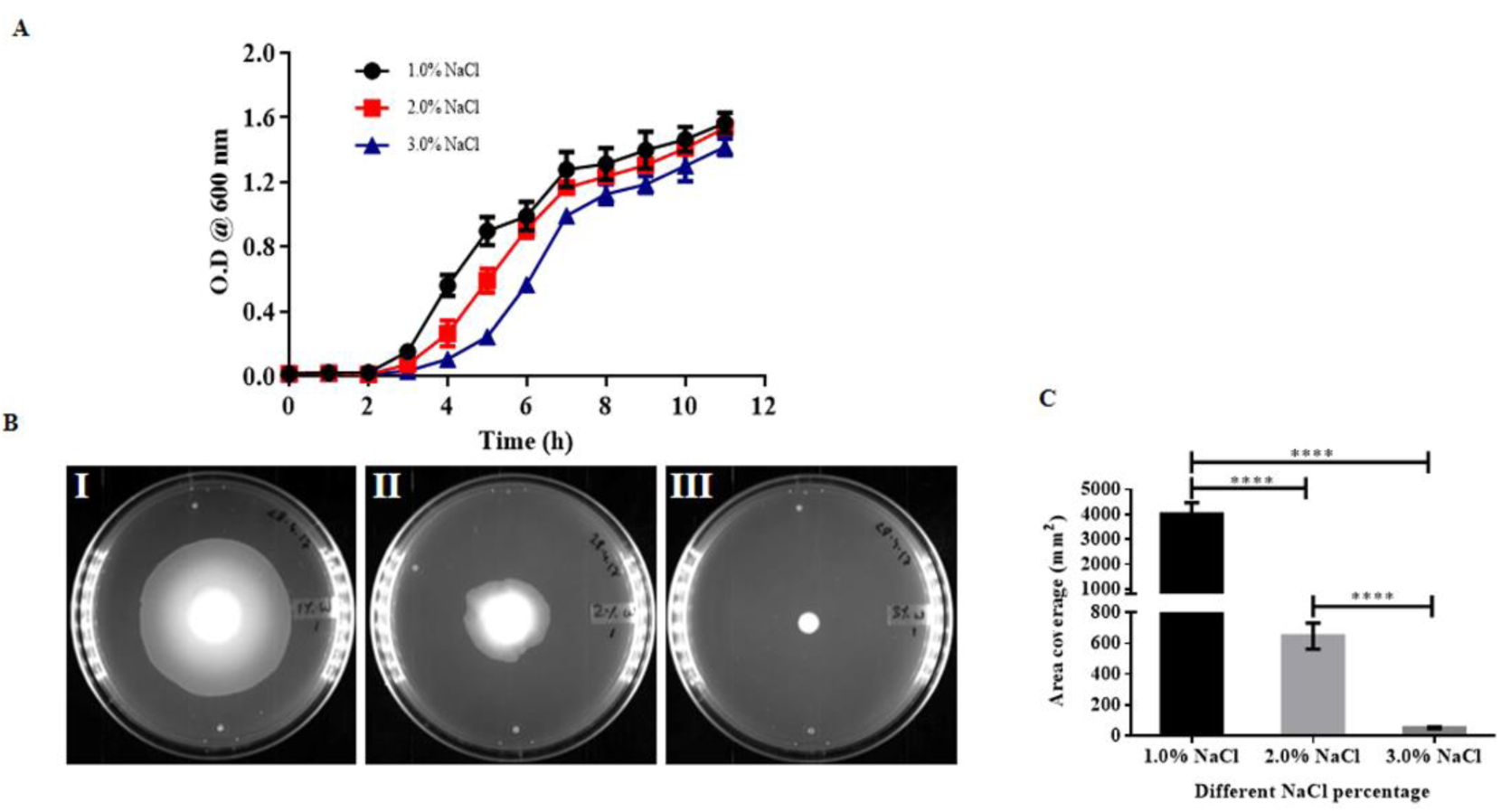
(A) Growth kinetics of *Bacillus cereus* MHS in Luria broth in the presence of different NaCl concentrations. (B) Swarming motility of *Bacillus cereus* MHS after 12h in the presence of (I) 1.0% NaCl (II) 2.0% NaCl and (III) 3.0% NaCl in 0.7% LA. (**C)** Area covered by *Bacillus cereus* MHS colonies in the presence of different NaCl concentrations after 12 h of growth: All results are shown as means ± S.E. (N=9). Statistical significance was analyzed using Student’s *t*-test (2-tailed, unpaired). *P-value* with less than 0.05 were considered to be statistically significant (\**P-value* ≤ 0.05; \*\**P-value* ≤ 0.01; \*\*\**P* value ≤ 0.001; ****, *P-value* ≤ 0.0001)

### 3.3. Alterations of arrangement, orientation and dimensions of B. cereus MHS cells during swarming on LA plates in the presence of different salt concentrations

CLSM is one of the most valuable research tools, which helps to study the cellular arrangement, orientation, morphology, and real-time visualization of fully hydrated and living bacteria in a colony. The *B.cereus* MHS cells were grown on 0.7% LA plates for 12h in the presence of varying NaCl concentrations (1.0, 2.0, and 3.0%) and bacterial cells at the periphery of the growing colonies were observed by CLSM. The CLSM images revealed that although a reduction in the growth of colonies occurred in the presence of increased salt concentrations in growth media on LA plates, the overall pattern of cell orientation and arrangement did not alter in different salt concentrations. The cells were observed to be arranged in random patterns in the presence of all the concentrations of NaCl **(Figure 3)**. However, the morphological parameters like length, width, and surface area to volume ratio of *B.cereus* MHS were observed to be altered in the presence of increased salt concentrations **(Figure 4)**. A gradual decrease in the length of the cells was observed, along with an increase in salt concentrations. In 1.0% NaCl, the length of MHS cells was 11.01 ± 0.18 μm while the lengths were observed to be 9.27 ± 0.15μm and 8.51 ± 0.17μm in the presence of 2.0% and 3.0% NaCl, respectively. Interestingly, the width of MHS cells did not appear to change significantly in between growth in the presence of 1.0 % (1.3 ± 0.006µm) and 3.0% NaCl (1.2 ± 0.006 µm) but it changed when compared between 1.0% and 2.0% NaCl (1.3 ± 0.008 µm). By considering cells as cylinders, the surface area to volume ratio of MHS cells was calculated and the ratio was found to be also altered in the presence of 2.0% and 3.0% NaCl as compared to colonies grown in the presence of 1.0% NaCl. The surface area to volume ratios were found to be 3.28 ± 0.017 μm^-1^, 3.47 ± 0.018 μm^-1^, and 3.37 ± 0.024 μm^-1^ in the presence of 1.0%, 2.0%, and 3.0% NaCl respectively.

**Fig. 3.**
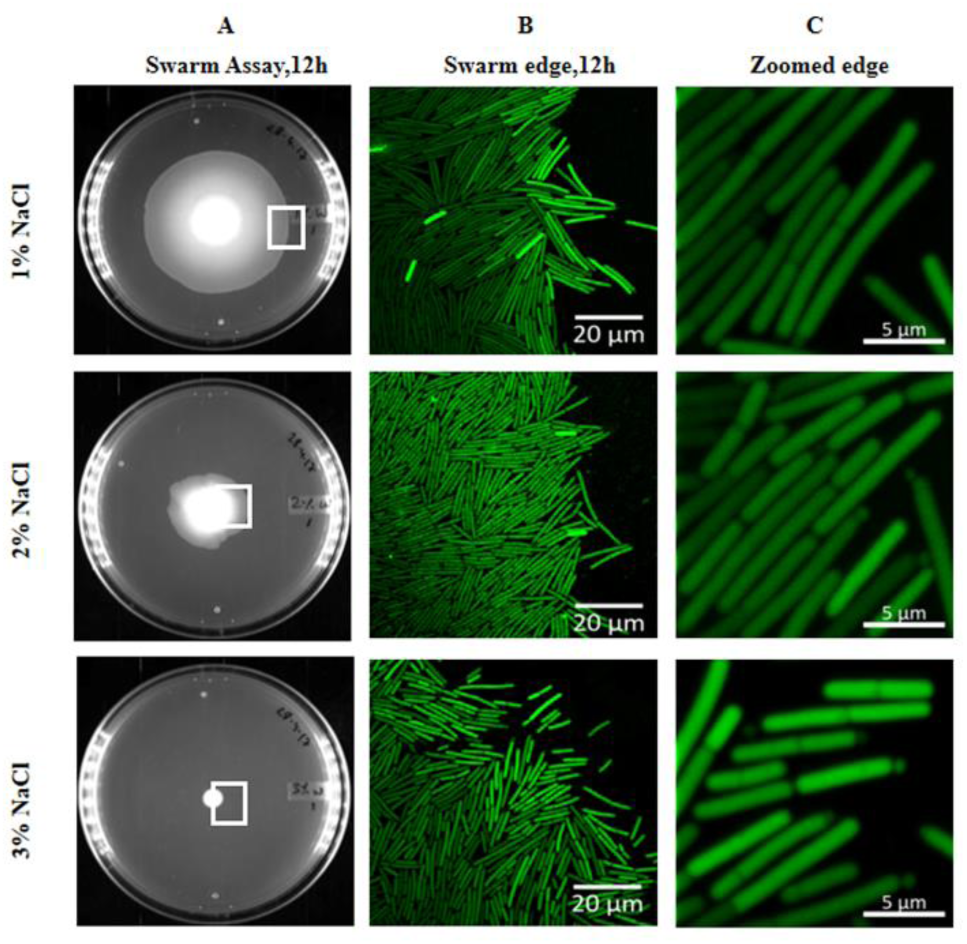
Representative images of *Bacillus cereus* MHS grown for 12 h in the presence of different concentrations of NaCl in 0.7% LA. (A) Swarming motility, (B) 100X CLSM images of peripheral cells of growing colonies, (C) Zoomed CLSM image of peripheral cells

**Fig. 4.**
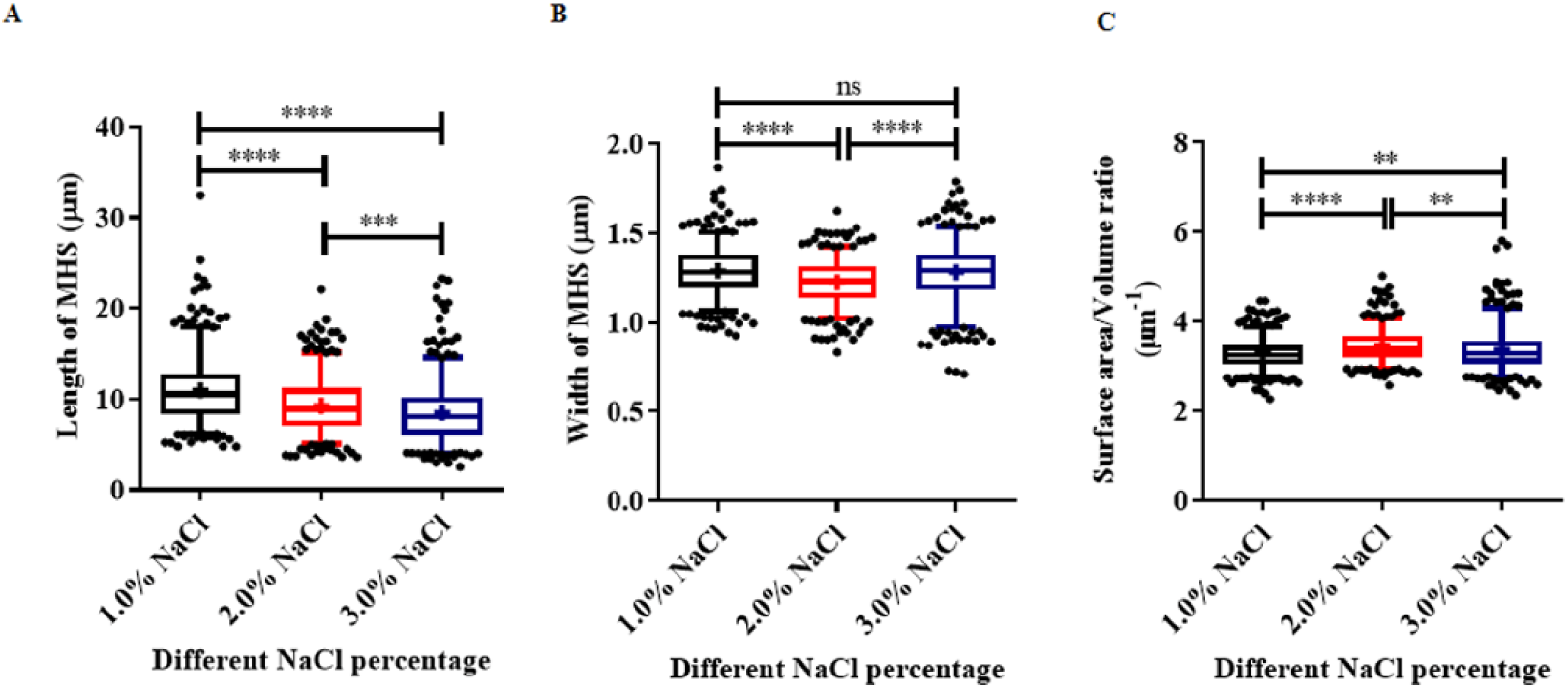
Morphological parameters of peripheral *Bacillus cereus* MHS cells grown under different NaCl concentrations in 0.7% LA. Box and whiskers plot of (A) Length in µm, (B) Width in µm, and (C) Surface area to volume ratio of *Bacillus cereus* MHS after 12 h (N=400), as calculated from CLSM images. Box plot crosses are the mean value. The boxes indicate the lower and upper quartiles and the central line indicates the median. Whiskers above and below the box indicate the 95th and 5th percentile. Cell volume was calculated as a cylinder using cell length and width

### 3.4. Effect of differential salt concentrations on flagellation of Bacillus cereus MHS

To observe the level of flagellation in *Bacillus cereus* MHS during swarming on 0.7% LA plates in the presence of different NaCl concentrations, transmission electron microscopy (TEM) was performed. The *Bacillus cereus* MHS strain revealed the presence of peritrichous flagella on the cell surface in the presence of 1.0% NaCl. Interestingly, with the increment of NaCl concentrations (from 1.0-3.0%), the number of flagella was found to be drastically decreased (**Figure 5**). This could be correlated to the observed decrease in swarming of MHS in the presence of increased NaCl concentrations.

**Fig. 5.**
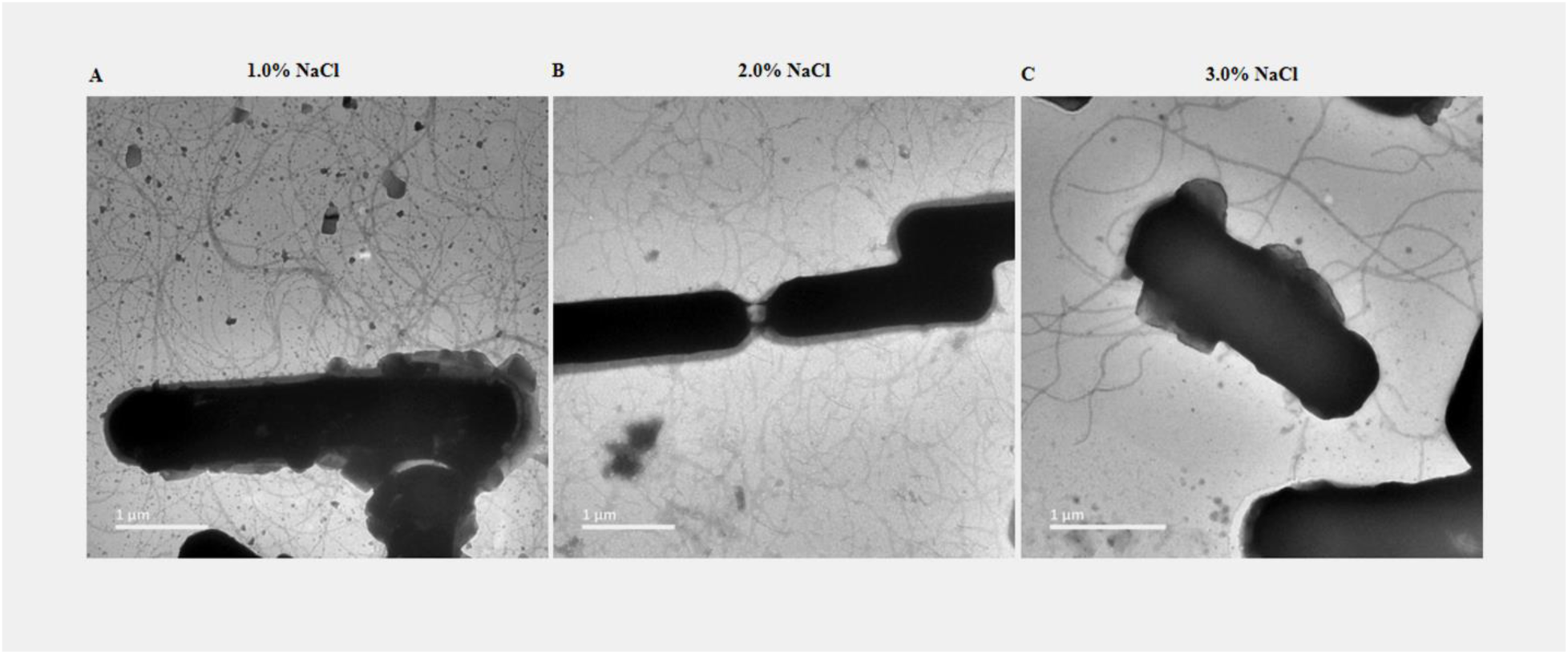
Representative images of TEM of peripheral cells of *Bacillus cereus* MHS after 12 h of growth in 0.7% LA in the presence of different NaCl concentrations (A) 1.0% NaCl (B) 2.0% NaCl and (C) 3.0% NaCl

### 3.5. Salt mediated change in surface topography of Bacillus cereus MHS

The surface topology and structural architecture of *Bacillus cereus* MHS during swarming on 0.7% LA under different NaCl concentrations were visualized using FESEM. The overall surface topographies of MHS cells were observed to be similar with different salt concentrations. Strikingly, we observed the presence of nanotube(s) connecting two or more cells along with nanotube networks and nanotube web in the growing colonies of MHS (**Figure 6A**).

**Fig. 6.**
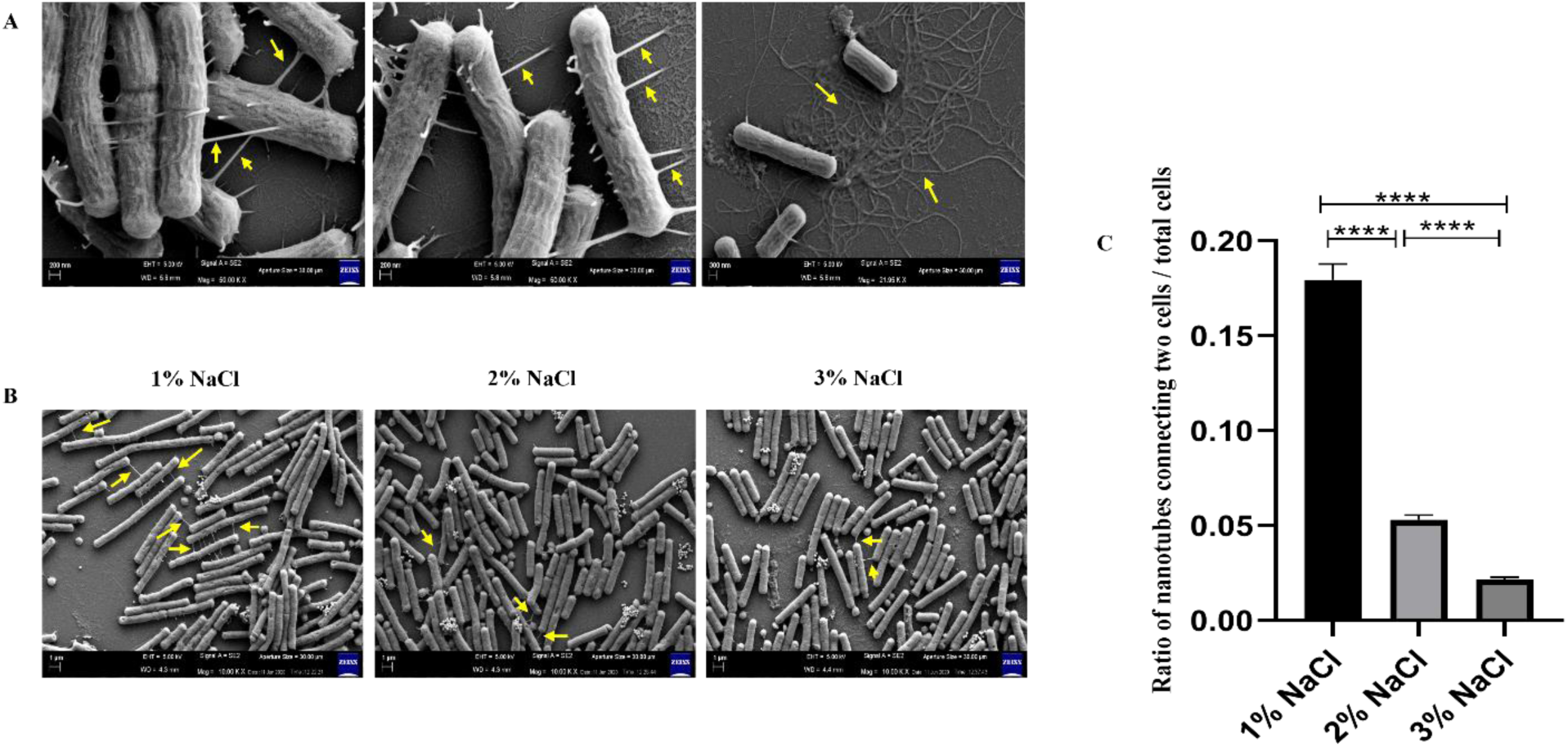
(A) Different types of bacterial nanotubes present in the peripheral cells of swarming *Bacillus cereus* MHS when grown in 0.7% LA (B) The presence of bacterial nanotubes in the peripheral cells of *Bacillus cereus* MHS grown in 0.7% LA with increasing NaCl concentrations (C) The ratio of two cells connected via nanotubes per total number of peripheral MHS cells

Although the presence of bacterial nanotubes was reported earlier in *Bacillus subtilis* (Dubey and Ben-Yehuda 2011), the present study revealed the presence of nanotubes for the first time in a *Bacillus cereus* strain. The average width of the nanotubes was measured and observed to be 69 ± 2.6 nm. Nanotubes of similar dimensions have been previously reported for other bacteria (Dubey et al. 2016; Patel et al. 2021). These observed structures differ from flagella or pili as the width of bacterial flagella falls within the range of 10 to 30 nm (Zhou & Li, 2015) and the width of *Bacillus cereus* pili is reported to be 6.8 nm (DesRosier & Lara, 1981).

Interestingly, it was found that the number of nanotubes, in the case of *B. cereus* MHS, varied with different salt concentrations which regulated the extent of swarming of MHS cells and the nanotubes were observed to be more in number in the presence of 1.0% NaCl as compared to 2.0 and 3.0% NaCl (**Figure 6B**). For estimation of the number of nanotubes, 15, 13, and 18 images from three separate experimental repeats were considered for 1-, 2-, and 3% NaCl, respectively. From each FESEM image, the ratio of nanotubes connecting two cells out of the total number of cells was calculated. Thereafter, to equalize the number of FESEM images in different concentrations of NaCl (1%, 2%, and 3%), a random selection of 10 such ratios was done ten times and average values for each selection were represented with SE (**Figure 6C**). As evident from **Figure 6C**, the average ratio of nanotube connecting two cells decreased with the increase in salt concentrations in the swarm media as 0.179±0.008 (1% NaCl), 0.053±0.002 (2% NaCl), 0.021±0.001 (3% NaCl). It was also observed that the number of nanotubes attached to the substratum also apparently reduced with increased NaCl concentrations during the swarming of MHS cells on semi-solid surfaces.

### 3.6. Differential expression of genes during swarming of MHS in the presence of salt

The differential expression of genes involved in the formation of flagella (*bfla),* the formation of nanotubes (*ymdb*), and regulating quorum sensing (*plcr),* was measured by semi-quantitative RT-PCR and the level of expressions of different genes was compared after normalizing with the expression of housekeeping gene (*gatB_Yqey*). The expression of *bfla* was found to be decreased by 0.82±0.055-fold in the presence of 2.0% NaCl and by 0.63±0.067-fold in the presence of 3.0% NaCl when compared to the 1.0% NaCl (**Figure 7**). This decrease in *bfla* expression in the presence of increased NaCl justifies the observed decrease in motility of natural hyperswarming MHS strain on increasing the NaCl percentages in semi-solid LA media. The expression of the *ymdb* gene was found to be decreased by 0.33±0.057 and 0.404±0.191 folds in the presence of 2.0% and 3% NaCl respectively supporting the observed decrease in nanotube formation during swarming under 2.0% and 3.0% NaCl as compared to 1.0% NaCl **(Figure 7)**. Interestingly, the expression of the quorum sensing regulator gene *plcr* was found to be decreased by 0.53±0.145-fold in the presence of 2.0% NaCl and by 0.43±0.119-fold in the presence of 3.0% NaCl when compared to 1.0% NaCl (**Figure 7**) which suggests that the hyperswarming motility in MHS is positively regulated by quorum sensing.

**Fig. 7.**
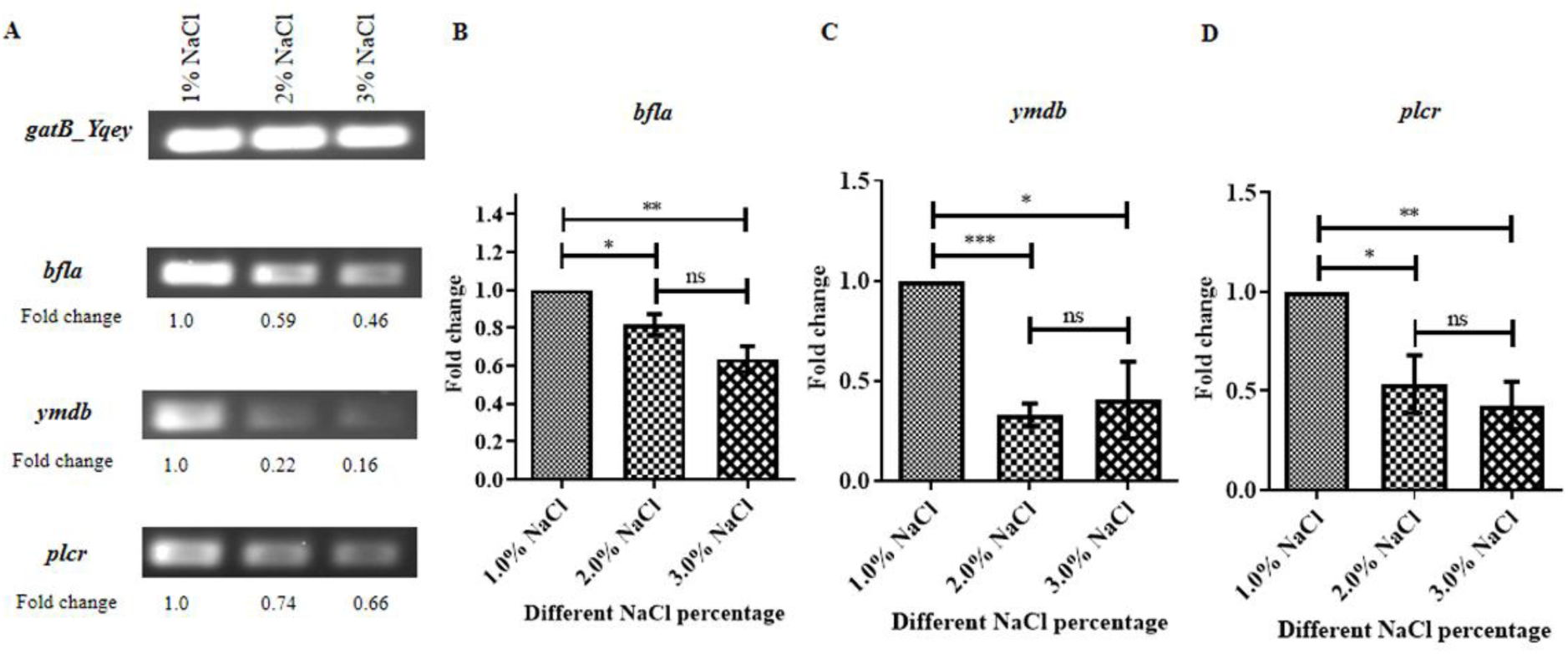
Differential expression of *bfla, ymdb,* and *plcr* genes in the peripheral cells of *Bacillus cereus* MHS colonies grown in the presence of varying concentrations of NaCl in 0.7% LA. (A) Representative images of semi-quantitative RT-PCR amplified DNA of respective genes in agarose gels (N = 3). Relative expressions of *bfla* (B), *ymdb* (C), and *plcr* (D) were normalized with the expression of *gatB_Yqey* gene (Housekeeping). For 1.0% NaCl conditions, the relative expressions were considered as 1

## 4. Discussion

An attempt was made to examine the hyperswarming behaviour of the *Bacillus cereus* MHS strain on 0.7% L.A medium at different salt (NaCl) concentrations. In terms of swarming motility and colony growth on semi-solid surfaces, the hyperswarmer (MHS) strain of *Bacillus cereus* responded by reducing its hyperswarming activity in the presence of increased salt concentration. Several other phenotypes and cellular ultra-structures were also altered in the presence of increased salt concentrations in growth media. Bacterial cells tend to change their morphological parameters to adjust to the external conditions and in the case of MHS such changes were to adapt to different osmolarity and associated stress during growth and as survival strategies. Since *Bacillus cereus* MHS could move faster, it could quickly change this property when an alteration of osmolarity in the surrounding environment was introduced.

With the help of CLSM studies, it was found that the morphological parameters like cell length, cell width, and surface area to volume ratio altered with increasing concentrations of NaCl. At the growing edges of MHS colonies, heterogenous distribution of *Bacillus cereus* MHS cells for length, width, and surface area/volume were observed suggesting that various sizes of cells were present at the peripheral colonies of *Bacillus cereus* MHS strain. Interestingly the randomness or heterogeneity in cell length increased with increased salt concentrations indicating unfavorable conditions were imposed by the increased amount of salt. However, the average length of cells was found to decrease with an increase in the NaCl concentrations. Since the hyperswarming of bacterial cells has been reported to be associated with cell elongation (Kearns 2010) hence the observed reduction in the cell length in the increased concentrations of NaCl can be correlated with the observed reduction in hyperswarming of *Bacillus cereus* MHS in this study. The width of the cells was found to be altered in the presence of increased NaCl concentrations. The randomness in width did not alter much with an increased amount of salt. Moreover, the SA/V ratio was also observed to be altered significantly. The different surface area to volume ratio of the *Bacillus cereus* MHS cells grown in the presence of different salt concentrations suggests that cells were adjusting their cell contents to adjust to the external changes in osmolarity when NaCl concentrations were varied.

Further, the transmission electron microscopic studies revealed that the salt-induced decrease in swarming motility was associated with decreased flagellation in the hyperswarmer *Bacillus cereus* MHS strain. The current finding corroborates with the findings of Murugan et al. (2024) where an increase in swarming motility was observed with increased flagellation at 2.0% of NaCl concentration. It thus implies that the presence of salt as a nutrient, as well as a mediator of osmolarity of the growth media, is an important factor in regulating the physiology and survival of bacteria.

The most interesting observation in the present study was the discovery of the presence of bacterial nanotubes in a naturally hyperswarming *Bacillus cereus* MHS strain. Through Scanning Electron Microscopy, nanotubes connecting the bacterial cells were observed. The bacterial nanotubes are hollow connecting structures between cells as well as substratum, which allows for the internal transport of various metabolites, and signaling molecules (Patel et al., 2021). Besides regulating the hyperswarming, nanotubes have been reported to be involved in the exchange of molecular information (both genetic and cytosolic) between and among bacteria (Ficht 2011, Baidya et al. 2018, Shitut et al. 2019, Baidya et al. 2020). The bacterial nanotubes have also been shown to be involved in cell death, with a possibility of altruistic behaviour of the neighbouring bacterial cell (Pospíšil et al., 2020). The nanotubes between interconnected bacterial cells can make bacteria act like multicellular entities, thus giving them ecological benefits. Such physical attachments help bacteria to establish a kind of ordered behaviour in the cells just like swarming. The interesting finding in the present study was that the number and overall network of bacterial nanotubes was reduced in MHS with the increase in NaCl concentrations indicating a possible link between bacterial nanotube formation and hyperswarming ability in *Bacillus cereus* MHS. The findings from the present study thus open new doors in understanding the role of bacterial nanotubes in bacterial coordinated movement (hyperswarming), survival, and pathogenicity.

The alterations in the cellular behaviours of bacteria occur due to altered expression of genes and proteins involved in maintaining cellular homeostasis to help the bacteria adapt to external environmental conditions. To understand the differential expression of genes involved in the hyperswarming of MHS under different NaCl concentrations, genes involved in quorum sensing (*plcr*), flagellation (*bfla*), and nanotube formation (*ymdb*) were explored.

*plcr*, the master regulator of quorum sensing (QS) regulates over 40 different genes that govern the ion balance, food supply, release of bacterial toxins as well as cell-cell interactions (Lereclus et al. 1996, Gohar et al. 2008) which helps in cell survival. The *plcr* regulon appears to integrate the external environmental signals (food deprivation, self-cell density) and regulates the transcription of its target genes to overcome the challenges posed to the bacteria which might hinder its growth, hence, *plcr* has also been referred to as key component in bacterial adaptation to its environment (Gohar et al. 2008). *plcr* has been shown to positively but indirectly regulate the flagellin expression in the *Bacillus cereus* ATCC 14579 strain and the mutant *plcr* decreased the flagellin production by 3 folds (Gohar et al. 2002) and the overall swarming motility in *Bacillus cereus* swarm cells (Senesi et al. 2010). In the contrary, a study conducted by Salvetti et al. (2011) showed that swarming in *B. cereus* was associated with a reduction in *plcr* expression. Interestingly, in the present study, it was observed that NaCl induced decrease in swarming and reduction in flagellation was associated with the decrease in *plcr* and *bfla* expression which indicates the involvement of *plcr* in the regulation of flagellation and swarming with a positive correlation in *Bacillus cereus* MHS cells which indicates that in most of the cases *plcr* regulates flagellation and swarming in *Bacillus cereus* with a positive correlation (Senesi et al. 2010). *ymdb* gene has been reported in *Bacillus subtilis* and shown to be involved in nanotube formation (Dubey and Ben-Yehuda 2011) and *ymdb* was also reported to be involved in late-adaptive response in a *Bacillus subtilis* strain (Diethmaier et al. 2014). A study conducted by Zhang et al. (2020) showed that the deletion of the *ymdb* gene helps in the swarming motility of *B. cereus* 0-9 strain. Contrary to this, results from this present study showed that the swarming motility and expression of the *ymdb* gene were reduced with increased NaCl concentrations in the semi-solid media. This could be because they are different *Bacillus cereus* strains and might have differences in regulatory networks. Another recent study reported that *ymdb* gene is present in many bacteria and its product can act as the stress diminishing factor for the bacteria to adapt to the changing environment (Zhang et al. 2020). *Bacillus cereus* MHS adapts itself to the increasing NaCl concentrations by decreasing the *ymdb* expression, which is supported by decreasing number of nanotubes. The research regarding the structure, roles, and consequences of bacterial nanotubes in ecological stress situations and swarming is very limited. Therefore, the findings from the present study pave the way for further investigations into the functions of nanotubes in multicellular behaviours in bacteria including *Bacillus* sp.

## 5. Conclusion

In conclusion, the present study revealed that the reduction in hyperswarming abilities of *Bacillus cereus* MHS in the presence of an increased amount of NaCl in the growth medium is mediated through decreased flagellation and decreased nanotube formation. The electron microscopic (SEM and TEM) observation of decreased numbers of flagella and nanotubes in the hyperswarming MHS strain of *Bacillus cereus* in the presence of increased salinity is supported by the results of differential expressions of *bfla* and *ymdb* genes. This signifies the involvement of flagella and nanotubes in hyperswarming motility in *Bacillus cereus* MHS. As the expression of QS regulator gene *plcr* was found to be reduced with NaCl induced decreased swarming of *Bacillus cereus* MHS strain, the hyperswarming motility in MHS is likely to be regulated by QS and represents a multicellular behaviour. The results of the present study open new avenues for exploring the roles of nanotubes in the multicellular behaviours of *Bacillus cereus*.

## 6. Acknowledgments

The authors thank Mr. Ritabrata Ghosh for Confocal imaging, Mr. Kasinath Sahu for Scanning Electron Microscopy, and Mr. Sumanto Moi for Transmission Electron Microscopy. NKB acknowledges UGC-JRF for research fellowship. KP acknowledges the Department of Biotechnology, Government of India (DBT-India) for providing a research fellowship. NG acknowledges IISER Kolkata for a postdoctoral fellowship.

## 7. Author Contributions

NKB and TKS conceived the study and designed the experiments. NKB performed the experiments. HS isolated the strain *Bacillus cereus* MHS. NKB, KP, NG and TKS analyzed the data and wrote the manuscript.

## 9. Availability of Data and Materials

All data generated or analyzed in this study are included in the article.

## 10. Declaration

### Ethical Approval

Not applicable.

### Consent to Participate

All authors had consented to participate in the study.

### Consent for Publication

All authors have given consent for publication.

