## Supplementary File for "Salt-induced reduction of hyperswarming motility in *Bacillus cereus* MHS is associated with reduced flagellation, reduced nanotube formation and reduction in expression of quorum sensing regulator"

### Figure S1

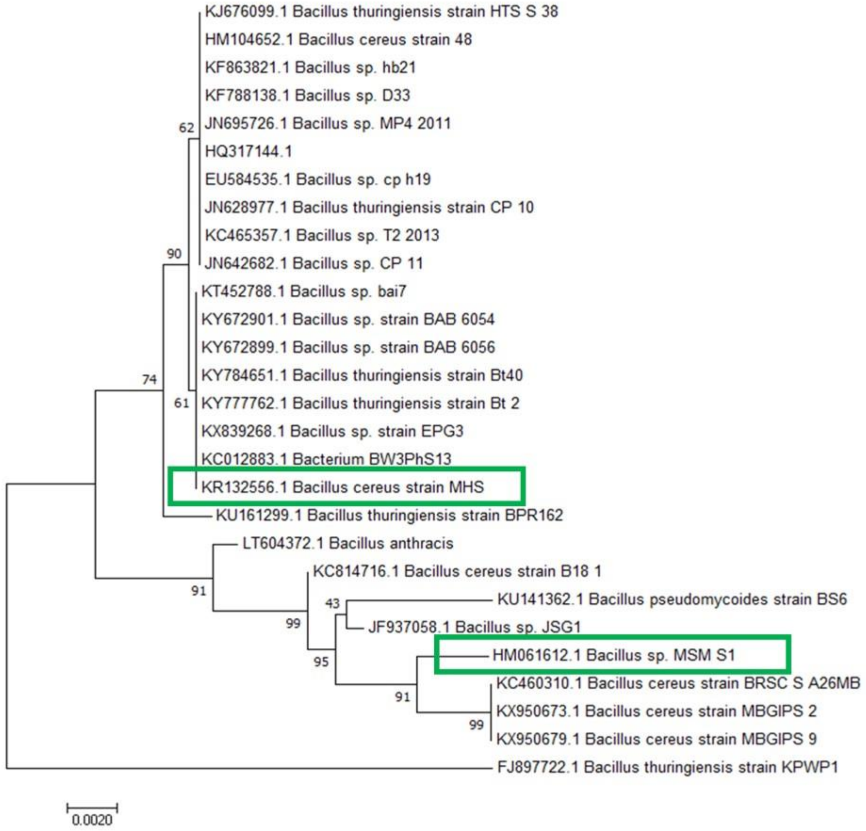


Figure S1. Phylogenetic relationship between *Bacillus cereus* MHS and MSM-S1 strains based on 16SrRNA gene sequence obtained by the Neighbor-joining (NJ) method. The number denotes the number of bases substituted per site. The accession number is presented before the species or strain name. Green box indicates the bacterial strain nomenclature box
